# MBIR: A Cryo-electron Tomography 3D Reconstruction Method that Effectively Minimizes Missing Wedge Artifacts and Restores Missing Information

**DOI:** 10.1101/355529

**Authors:** Rui Yan, Singanallur V. Venkatakrishnan, Jun Liu, Charles A. Bouman, Wen Jiang

## Abstract

Cryo-Electron Tomography (cryo-ET) has become an essential technique in revealing cellular and macromolecular assembly structures in their native states. However, due to radiation damage and the limited tilt range, cryo-ET suffers from low contrast and missing wedge artifacts, which limits the tomograms to low resolution and hinders further biological interpretation. In this study, we applied the Model-Based Iterative Reconstruction (MBIR) method to obtain tomographic 3D reconstructions of experimental cryo-ET datasets and demonstrated the advantages of MBIR in contrast improvement, missing wedge artifacts reduction, and missing information restoration compared with other reconstruction approaches. Considering the outstanding reconstruction quality, MBIR has a great potential in the determination of high resolution biological structures with cryo-ET.

## Introduction

Cryo-electron tomography (cryo-ET) has emerged as a promising technique that allows us to comprehensively explore macromolecular complexes and cellular architecture in near-native states ^1^. Using cryo-ET, the 3D tomogram of the biological sample can be reconstructed from a 2D tilt series collected by sequentially tilting the sample at different projection angles around a tilt axis ^2^. In practice, the quality of reconstruction with cryo-ET remains limited by several challenges in the data acquisition and reconstruction process.

The extremely poor signal-to-noise ratio (SNR) of cryo-ET is the first major challenge in improving cryo-ET resolution ^3^. To prevent significant radiation damage to biological samples by the electron beam, the total dose used for a cryo-ET tilt series is typically less than 100 e/Å^2^. This low-dose imaging strategy in combination with the increment of sample thickness during tilting results in very noisy, low contrast 2D projections, which poses a challenge in subsequent 2D tilt series alignments and deteriorates the resolution of cryo-ET 3D reconstruction ^4,5^.

The second major challenge of cryo-ET is the missing wedge artifacts caused by the limited tilt angle range during data collection ^6^. Since more electrons are lost to inelastic scattering as the effective sample thickness increases when the sample is tilted ^3^, the maximal tilt range of cryo-ET is typically restricted within ±70° to ensure enough electrons can traverse through the sample, generate elastic scattering, and form reliable images ^7^. Consequently, the absence of the high tilt angles (−90° ~ -70° and +70 ~ +90°) becomes a “missing wedge” of un-sampled information in Fourier space, leading to severe ray artifacts, structural elongation, and distortion effects in the final reconstruction ^8^. The missing wedge artifacts dramatically weaken the interpretability of the reconstructed tomogram and limit the achievable resolution of cryo-ET ^1^.

To address these challenges of cryo-ET, we introduce the Model-Based Iterative Reconstruction (MBIR) method ^9^ for tomographic reconstruction and benchmark the tomogram quality with the state-of-the art algorithms, including Back Projection (BP), Simultaneous Iterative Reconstruction Technique (SIRT), and Iterative Compressed-sensing Optimized Nonuniform fast Fourier transform reconstruction (ICON) ^10^. In MBIR framework, the reconstruction is formulated as the maximum a posterior (MAP) estimate of the unknowns, given the measurements

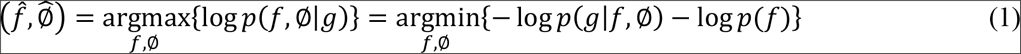

where *g* represents the data obtained from an imaging system (e.g. cryo-ET tilt series), *f* represents the unknown 3D structure to be discovered, ø represents the unknown nuisance parameters of the system such as beam intensity fluctuations and noise characteristics. *p*(*g*|*f*, ø) is the likelihood function that models how the observations are related to the unknowns, *p*(*f*) is the assumed prior distribution of the unknown structure. Here *p*(*g*|*f*, ø) and *p*(*f*) indicate the forward model of image formation and prior model of the tomogram in MBIR algorithm, respectively ^9^. Currently, the forward model accounts for the decay of electron beam intensity following Beer-Lambert Law and combines with the estimation of detector noise. The prior model uses a Gaussian Markov Random Field to account for diffuse or sharp interfaces between structural features and encourage smoothness in the solution. The goal of MBIR will be to compute a final estimate 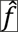 that represents a balance between fitting the measurements based on the system forward model *p*(*g*|*f*, ø) while remaining consistent with the prior model *p*(*f*). Fig. 1 illustrates a general framework of MBIR for solving inverse problems in imaging applications.

**Fig. 1.**
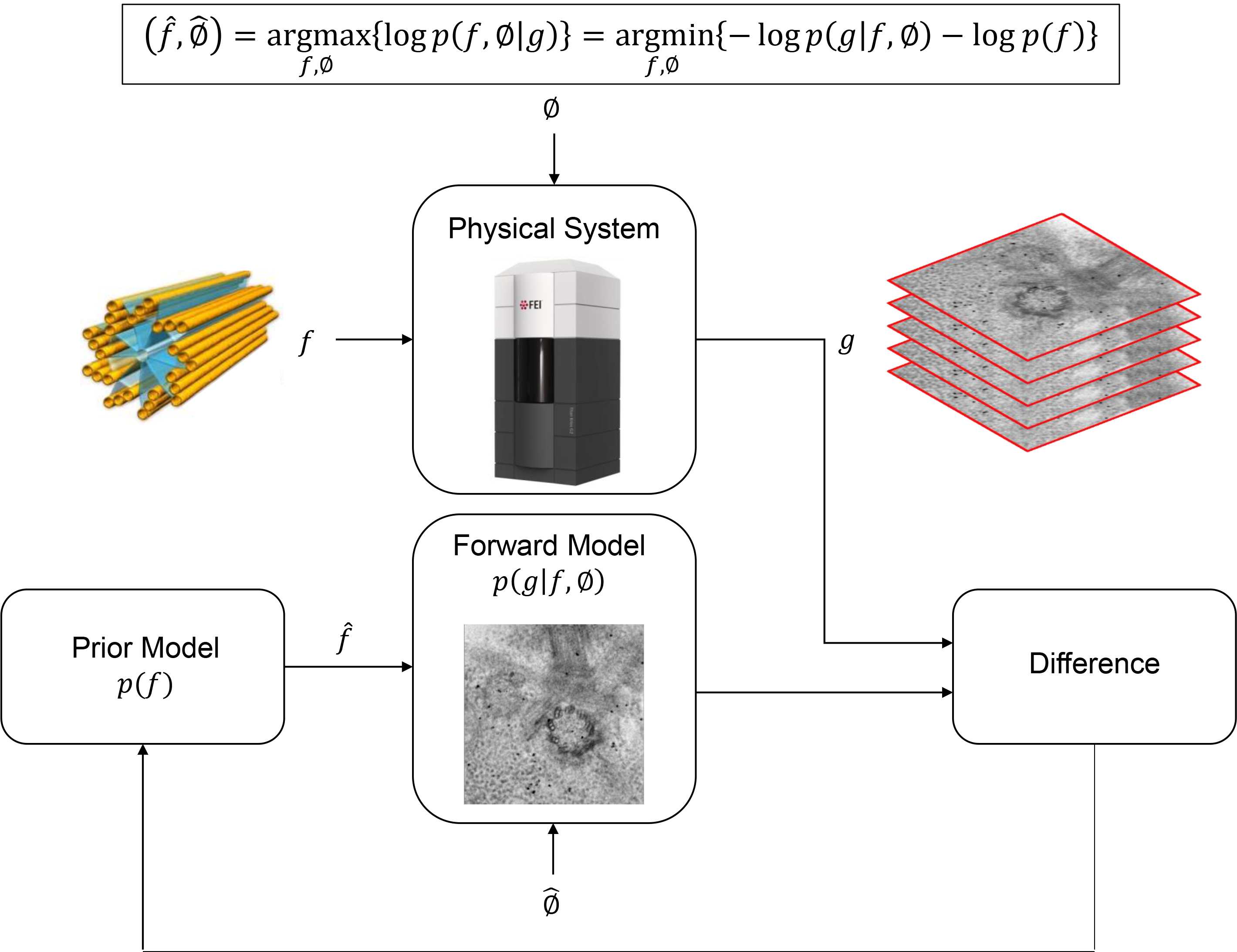
Graphical scheme of the MBIR algorithm. *g* denotes the tilt series from cryo-ET, *f* denotes the unknown structure, and ø denotes unknown nuisance parameters ofthe system (e.g. noise characteristics) which needs to be determined in the inverse process. *p*(.) denotes the probability density function and *p*(*g*|*f*, ø) and *p*(*f*) are the forward model and prior model in the MBIR algorithm, respectively. 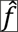 and 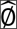 denote the estimate of *f* and ø, respectively.

MBIR method has been previously shown to generate better quality tomograms when applied to tomography applications like CT scan, X-Ray tomography, positron emission tomography (PET), optical diffusion tomography (ODT), and atomic resolution electron tomography of radiation-resistant material specimens ^11^. MBIR combines a forward model for image formation with a prior model for the unknown structure to reconstruct tomograms ^9^. In this study, tests with both plastic embedded ET dataset and ice embedded cryo-ET datasets have shown that MBIR can significantly improve the reconstruction quality with enhanced contrast, reduced missing wedge artifacts, and partially restored information in the un-sampled angular region.

## Results

### Missing wedge assessments using gold markers

We first evaluated MBIR using one cryo-ET dataset (EMPIAR-10045) by visually examining the missing wedge artifacts of gold markers in different slice views of the tomograms. Due to the missing wedge problem, the gold markers become elongated along the direction of the missing wedge and suffer from halos and streaking artifacts in the adjacent region. Fig. 2 compares slice views of the reconstructions generated by the four methods using gold markers as an indicator of quality. In each block, three planes represent the XY-slice (middle plane), XZ-slice (top plane) and YZ-slice (right plane) of the tomogram, respectively, intersecting at the same gold marker. The zoomed-in view of the gold markers pointed by white arrows in the three planes are placed at the corner of the corresponding planes. From the XY-slices of tomograms, it is clear MBIR (XY-slice in Fig. 2d) has eliminated the halos artifacts and displays more round, sharp-edged gold markers than other methods. In the XZ and YZ-slices, MBIR (Fig. 2d) significantly reduced the elongation and ray artifacts of gold markers with improved contrast of the biological structures, compared with the tomograms reconstructed by other methods. Hence, MBIR-reconstructed tomograms show less artifacts from the missing wedge problem, better contrast in cryo specimen, and clearer background.

**Fig. 2.**
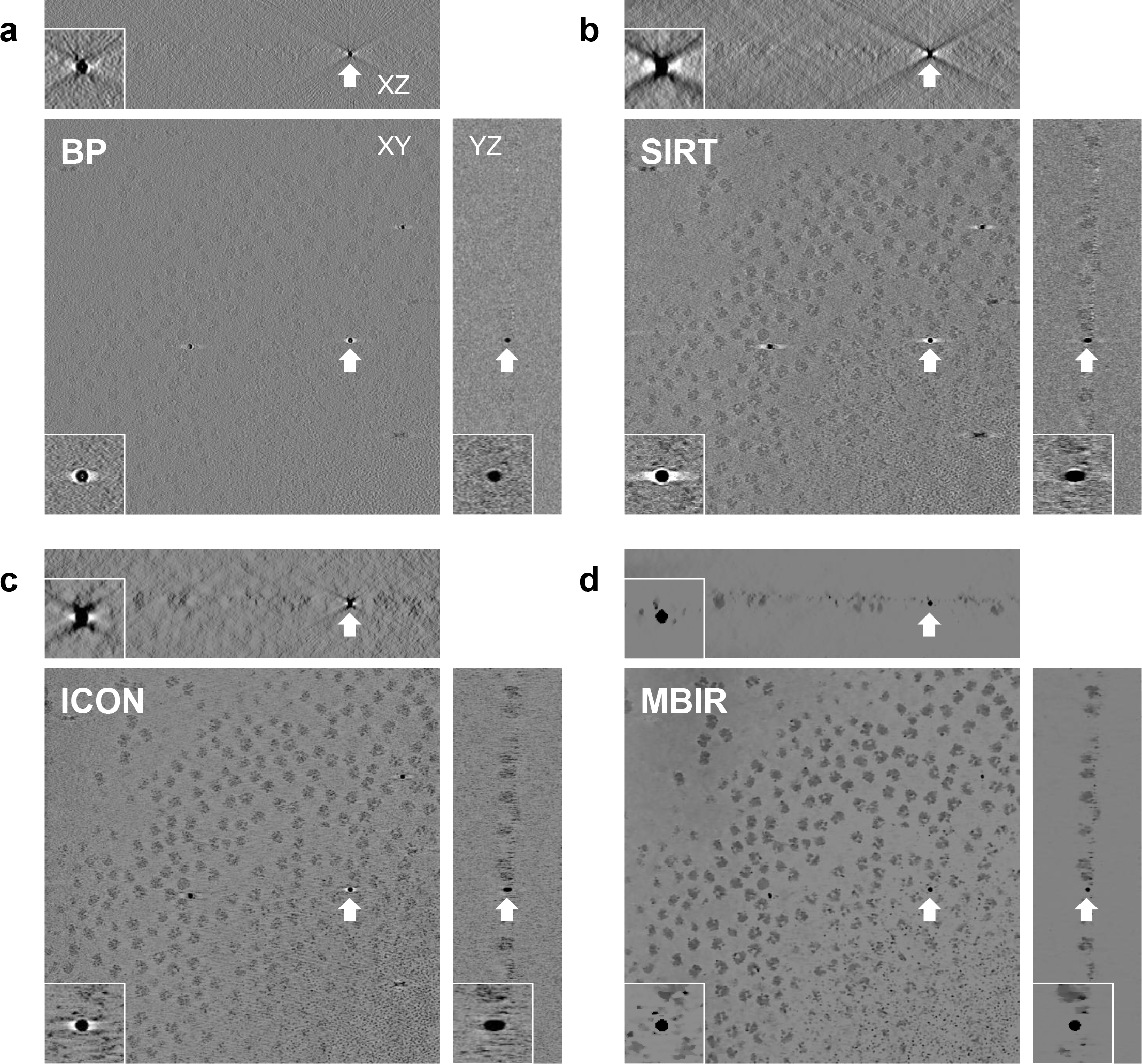
Comparison of tomograms from an experimental cryo-ET dataset (EMPIAR-10045) reconstructed by BP(a), SIRT (b), ICON (c) and MBIR (d) methods. The three planes for each method represent the XY-slice (middle plane), XZ-slice (top plane) and YZ-slice (right plane) of the tomogram intersecting at the same gold marker. In each plane, the gold marker is indicted by a white arrow with corresponding zoomed-in view showing the missing wedge artifacts.

To further examine the performance of MBIR, we applied it to one cryo-ET dataset acquired with VPP (EMPIAR-10064 in Fig. 3a), two cryo-ET datasets without VPP (EMPIAR-10037 and EMPIAR-10110 in Fig. 3b and c), and one plastic embedded ET dataset (IMOD tutorial dataset in Fig. 3d). Fig. 3 shows the slice views of these four datasets in which each row represents the results of one dataset reconstructed by the four methods and each column represents the results of one method applied to different datasets. In Fig. 3b and d, XY-slices are mainly used to reveal the reconstruction quality of sample areas without targeting at a gold marker because the sample and markers are not on the same XY plane. For a challenging dataset shown in Fig. 3b, it is clear that BP reconstruction quality is too poor to make the biological sample visible. SIRT and ICON reconstructions contain phantoms of gold markers at the upper left corners in XY-slice (circled by dash lines in Fig. 3b) which is caused by the missing wedge artifacts and should not appear here since gold markers are located in different Z sections of the sample. In stark contrast, MBIR in Fig. 3b is able to drastically reduce the missing wedge problem in XZ-slice and YZ-slice, completely suppress the gold marker phantoms in XY-slice and considerably enhance the contrast of biological samples. In addition, MBIR provides better quality of tomogram in other datasets of Fig. 3, which is in a good agreement with the results shown in Fig. 2. In summary, the comparison of slice views among different methods in Fig. 3 and Fig. 2 gives a clear impression that MBIR has superior performance in boosting contrast of biological specimens, eliminating halos and streaking artifacts, retaining sharp features, and reducing noise. The superior performance of MBIR is evident in both cryo-ET (Fig. 2 and Fig. 3a-c) and plastic-embedded ET (Fig. 3d) datasets.

**Fig. 3.**
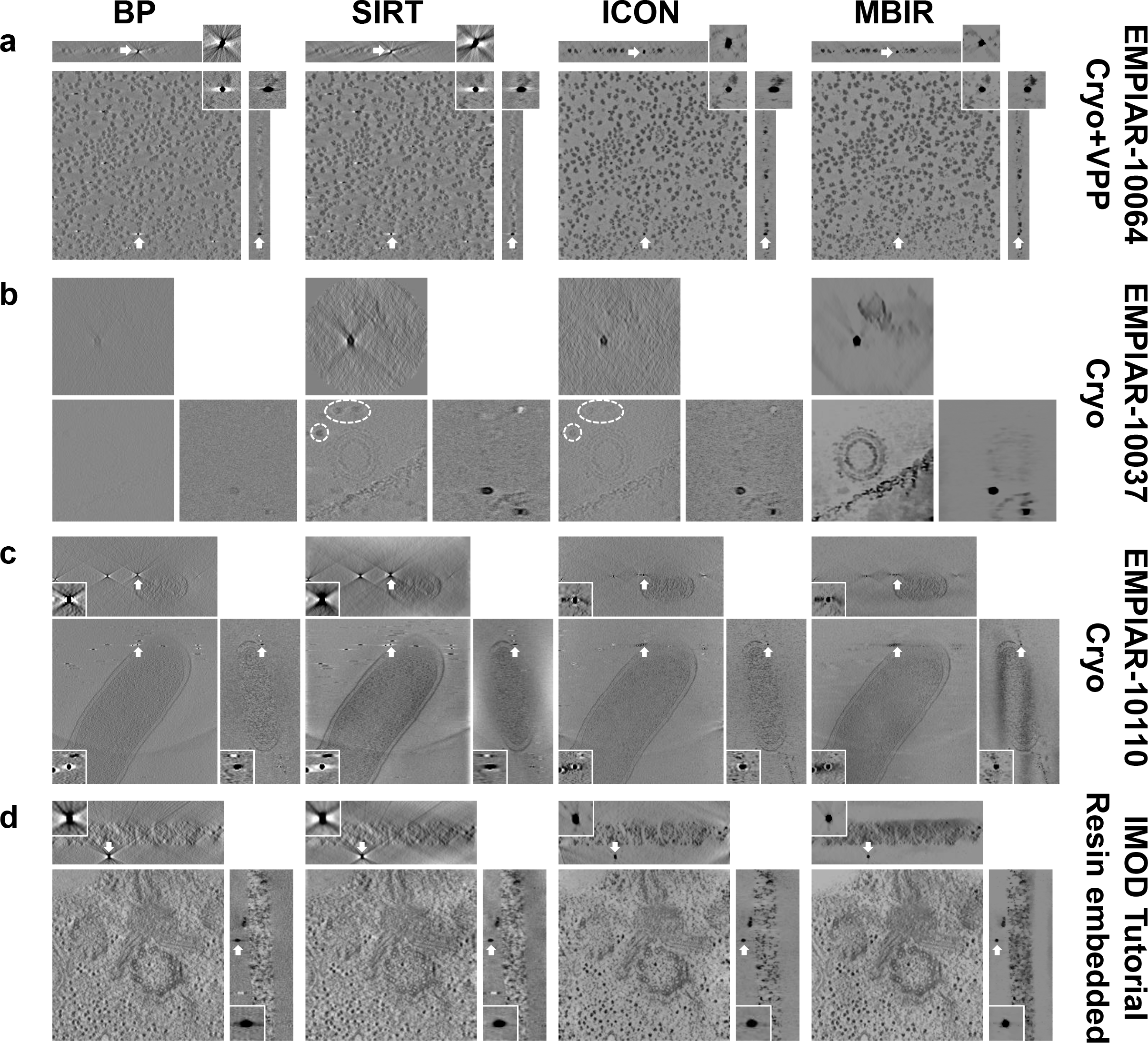
Comparison of tomograms from multiple experimental ET datasets reconstructed by different reconstruction techniques. Each row indicates the reconstructions from the same dataset using different methods. Each column indicates the reconstructions from the same method applied to different datasets. The data type and EMPIAR ID are denoted at the right side of each row. The method of comparison in each dataset is the same as described in Fig. 2. Note that the XY-slices of the dataset shown in (b) and (d) are used to show the biological sample area and not targeted at the gold markers since the sample and gold markers are located in different Z sections.

### Power spectra evaluation

To quantitatively evaluate MBIR’s ability in restoring missing information, we calculated the log-scaled power spectrum of the central XZ-slice and used it as a measurement of information restoration in 3D reconstruction. As depicted in Fig. 4, four plots of power spectra correspond to the central XZ-slices of the tomograms reconstructed by the four methods shown in Fig. 2. It is noted that MBIR can fill more un-sampled region in Fourier space than other methods, not only in the region of the missing wedge but also the empty space between two adjacent tilts, suggesting better performance of MBIR in restoring missing information. It is worth noting that the lines at the corners of BP (Fig. 4a) and SIRT (Fig. 4b) power spectra are due to the aliasing issue. To check if such aliasing issues are unique to our results, we downloaded another four 3D tomograms from EMDB ^12^, calculated their central XZ-slices power spectra, and plotted them in Supplementary Fig. 1. The results in Supplementary Fig. 1 suggest that this aliasing issue is a general phenomenon in the cryo-ET field since it is observed in a variety of data, including data from multiple research groups, varying TEM facilities and imaging conditions, a diverse range of samples, and different reconstruction software.

**Fig. 4.**
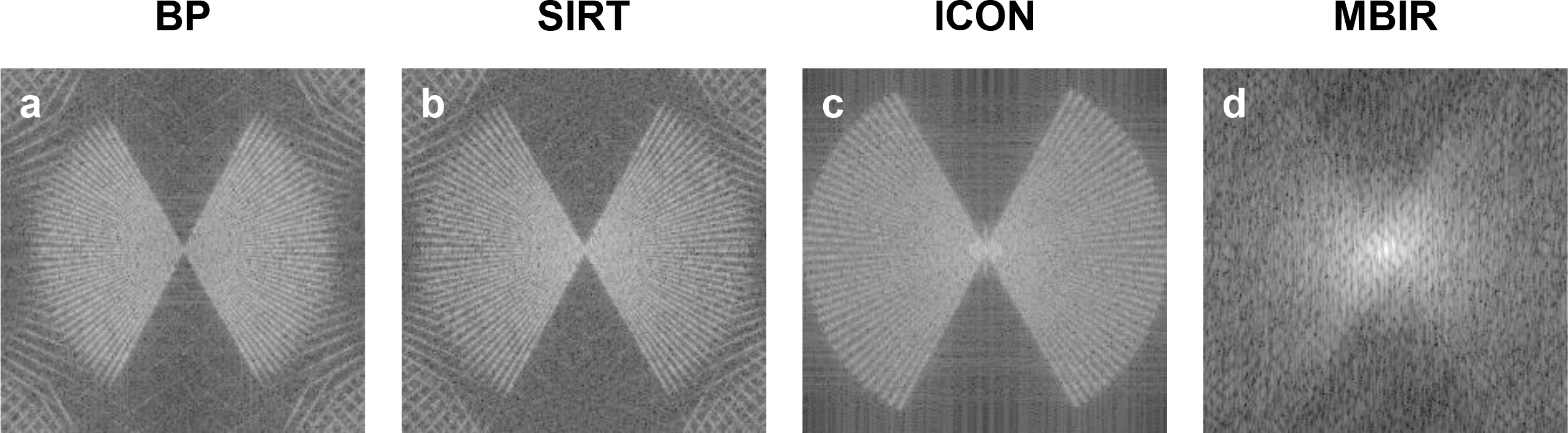
Comparison of the central XZ-slice power spectra from the tomograms shown in Fig. 2. The tomograms were reconstructed by BP (a), SIRT (b), ICON (c) and MBIR (d) methods, respectively.

We next examined the central XZ-slice power spectra of the datasets displayed in Fig. 3 and compared them in Supplementary Fig. 2. In general, MBIR and ICON yield more non-zero values in the missing wedge region than BP and SIRT, except for one challenging dataset (Supplementary Fig. 2b). However, power spectrum may not be a reliable and complete assessment for the information restoration because it only conveys the amplitude information without considering the phase information. What’s more, varying filters can be internally applied to tomograms in different methods to balance the non-uniform sampling in Fourier space ^13^. As a result, further validation is still needed to confirm the advantage of MBIR in restoring not only amplitude but also phase information.

### Cross validation of projections using the leave-one-out FRC method

We used the leave-one-out Fourier ring correlation (FRC) method ^14^ to explore the correctness of the information restored by MBIR and compare it with the performance of other reconstruction methods. In this test, the FRC is calculated for the raw tilt image *X* and the corresponding reprojection 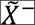 from a tomogram computed from all other tilts without tilt *X*. Here the tilde sign represents the reprojection from a tomogram, and the minus sign represents the tomogram used for reprojection is calculated by omitting the tilt *X* from the original tilt series to avoid bias. We first excluded a raw image *X* at a certain tilt angle and utilized the remaining images of the tilt series to generate a tomogram. Next, we re-projected this tomogram at the angle of tilt *X* to obtain a reprojection 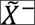. Finally, we calculated the FRC curve between the excluded raw image *X* and the reprojection 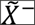, and used this FRC curve as a quantitative evaluation of phase information recovery. As shown in Fig. 5a, the first row and the second row are the raw images *X* (the first image in each row) and the reprojections 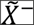 corresponding to different reconstruction methods at a smaller tilt angle 0° and a larger tilt angle 45°, respectively. The gold markers indicated by white arrows are zoomed in and placed at the lower left corners of each images. It is evident that the gold marker in MBIR reprojection is circular without discernible distortion or blurring, which is nearly identical to the original tilted image, even at a high tilt angle. In contrast, the gold markers in the reprojections of other methods clearly suffer from missing wedge artifacts including elongation, white halos, and blurring. Furthermore, such visual assessments are verified quantitatively by the FRC (Fig. 5b and c) of the raw tilt images and reprojections shown in Fig. 5a. As shown in Fig. 5b and c, the quick drop of BP (blue curve), SIRT (red curve) and ICON (green curve) FRC curves implies that only low resolution information is reliably restored in the non-sampled angular regions. However, the FRC curve of MBIR exhibits a significantly higher correlation between the reprojection and the original tilt image, confirming the successful restoration of the missing information.

**Fig. 5.**
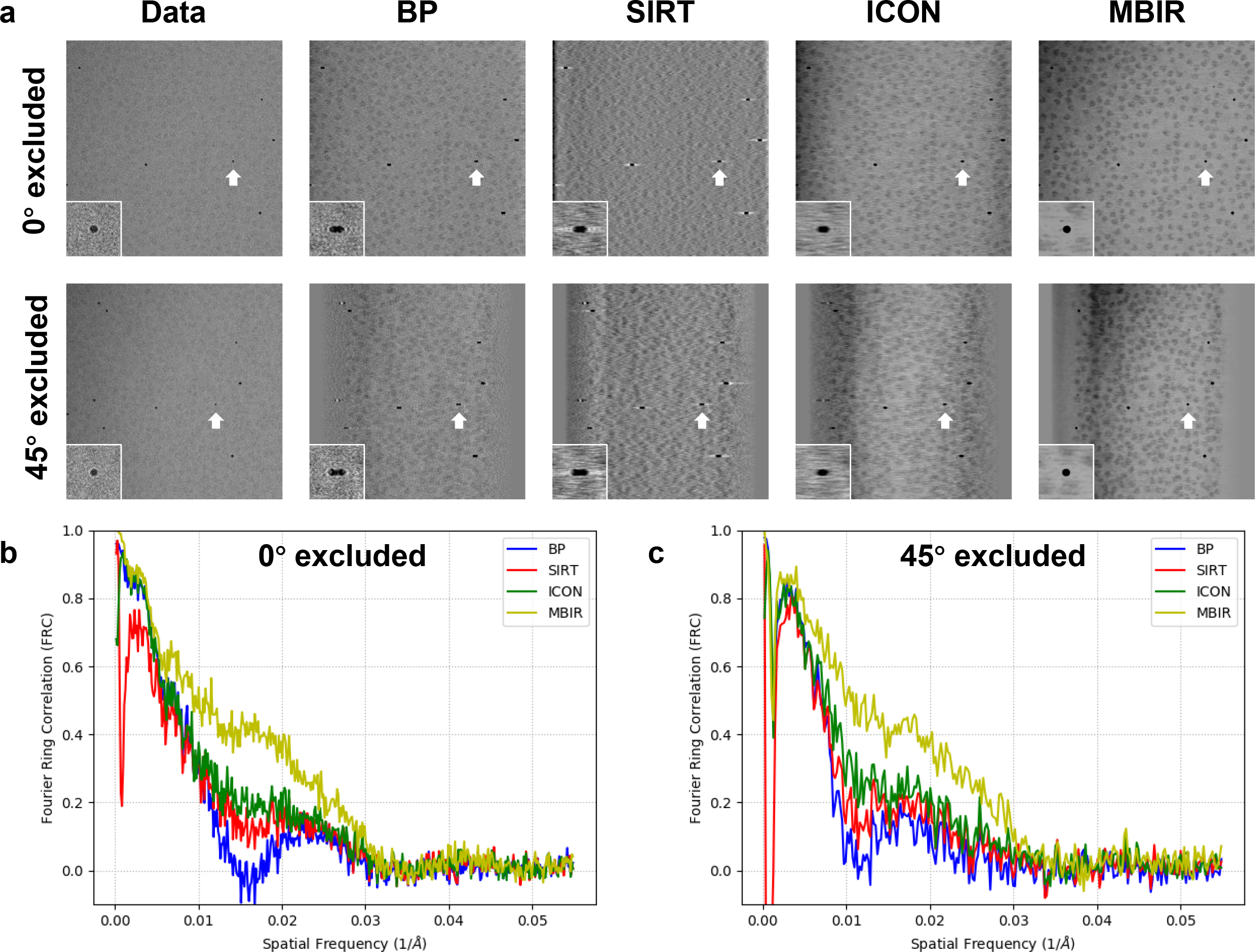
Comparison of missing information restoration from an experimental cryo-ET data (EMPIAR-10045) reconstructed by different reconstruction techniques using the leave-one-out FRC method. (a) Comparison of reprojections at two tilt angles (0° in the first row and 45° in the second row) using the tomograms generated without the corresponding tilt. The images in the first column are extracted from the tilt series, serving as the ground truth for comparison. In each plane, the gold marker indicted by a white arrow is displayed with corresponding zoomed-in view. (b) and (c) are comparisons of the FRC curves of reprojections against the ground truth as depicted in (a) when 0° and 45° tilt is excluded in the leave-one-out test, respectively.

To further substantiate the capability of MBIR in restoring missing information, we performed the same analysis as described in Fig. 5 on more datasets and summarized the comparisons of raw images and reprojections in Supplementary Fig. 3–4 and FRC comparisons in Fig. 6. As can be seen from Supplementary Fig. 3–4, MBIR preserved the round shape of gold markers in the leave-one-out reprojections at low (Supplementary Fig. 3) and high (Supplementary Fig. 4) tilt angles in both cryo (Supplementary Fig. 3a-c, Supplementary Fig. 4a-c) and plastic embedded datasets (Supplementary Fig. 3d, Supplementary Fig. 4d), which is consistent with the results shown in Fig. 5a. Fig. 6 shows the FRC comparisons of different methods when 0° (Fig. 6a, c, e, g) and 45° (Fig. 6b, d, f, h) tilts were excluded in the leave-one-out tests, respectively. The FRC curve of MBIR (yellow curve) in Fig. 6 is typically higher than that of other methods, which suggests the superior quality of MBIR in recovering authentic information of biological samples in 3D tomographic reconstructions. As demonstrated in Fig. 6a-b, VPP used in this dataset boosts the signal-to-noise ratio of cryo-ET images and improves the low frequency signal in FRC curve compared with Fig. 5b-c, leading to a smaller difference among the results of the four reconstruction methods than the case shown in Fig. 5. However, the local missing wedge artifacts around the gold markers remained in the tomograms reconstructed by the other three methods but not by MBIR for this cryo-ET dataset with VPP as shown in the corresponding slice views (Fig. 3a) and reprojections (Supplementary Fig. 3a and Supplementary Fig. 4a), emphasizing the advantages of MBIR method. Therefore, all the analyses above validate MBIR’s capability to partially restore the missing information in both cryo-ET and plastic embedded datasets.

**Fig. 6.**
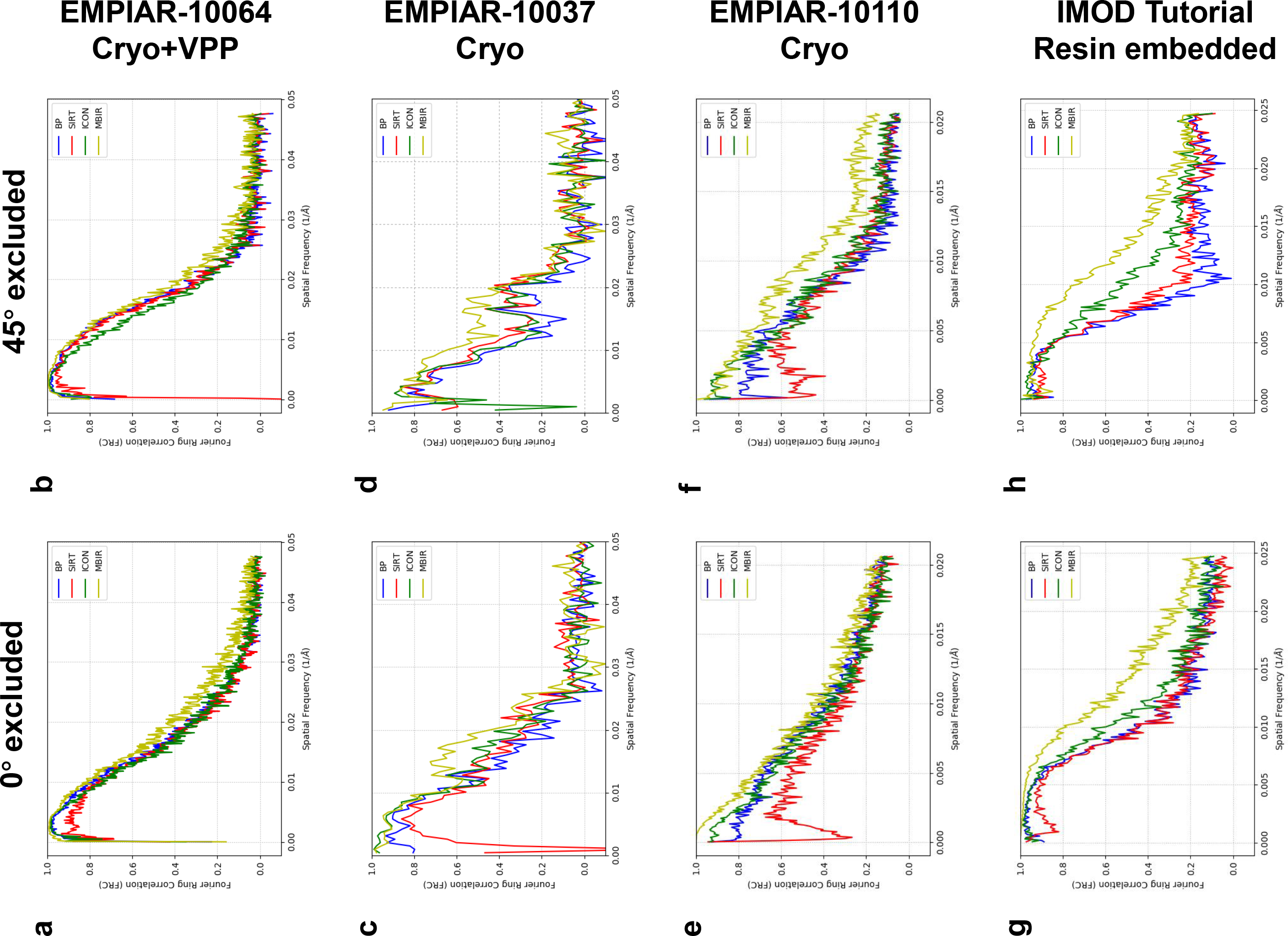
Comparisons of FRC curves from multiple experimental ET datasets reconstructed by different reconstruction techniques using the leave-one-out FRC method. Each row represents the comparison of FRC curves from the same dataset when 0° (left plot) and 45° (right plot) tilt is excluded in the leave-one-out test, respectively. The details of the corresponding reprojections and ground truths are shown in Supplementary Fig. 3 (0° excluded) and Supplementary Fig. 4 (45° excluded), respectively.

## Discussion

As a widely explored 2D/3D reconstruction method, MBIR has a growing impact on the medical, industrial, and scientific imaging fields. In the present work, we introduced the MBIR method into biological ET and corroborated the substantial advantages of MBIR over current, state-of-the-art reconstruction methods for both cryo and plastic embedded data. MBIR employs a model of the image formation process and combines it with a prior model of the 3D object to formulate a MAP estimation cost function which rejects measurements that do not fit the model. MBIR is finding a fit that balances between generating a reconstruction that matches the data and constraining it to have some properties that any real world object would have. Results on experimental data have effectively demonstrated the excellent performance of MBIR in contrast enhancement, missing wedge artifacts reduction, and missing information restoration, generating visually and quantitatively accurate tomograms.

Cryo-ET tomographic reconstruction usually suffers from problems such as high level of noise, poor contrast, artifacts caused by the missing wedge issue and unreliable restoration of missing information, which poses significant challenges to subsequent analysis of the tomograms. The clear benefits of MBIR should not only help achieve better quality reconstruction as shown in this work, but also facilitate further visualization and computational tasks, such as biological feature interpretation, structure segmentation, subtomogram averaging, and ultimately help advance cryo-ET to higher resolution.

While MBIR significantly improves tomography quality, the extensive computational load makes its speed slower compared to other approaches (Supplementary Table 1) and restricts the application of MBIR to large datasets. Recently, a computationally optimized algorithm termed Non-Uniform Parallel Super-Voxel (NU-PSV) has been developed for MBIR 3D reconstruction of CT images which enables rapid and massively parallel reconstruction while ensuring fast convergence ^15^. Thus, it is desirable to implement this powerful parallel algorithm into cryo-ET MBIR reconstruction in the future, using either GPU or multicore CPUs on multiple computer nodes. Furthermore, MBIR should be generalized to support tomographic reconstruction using double-tilt geometry and incorporate the objective lens contrast transfer function (e.g. defocus, astigmatism, Volta phase shift) into its forward image formation model during its iterative reconstruction process.

## Methods

### Implementation of MBIR

The MBIR algorithm was implemented as a standalone program using the C++ language by Dr. Charles Bouman’s group at Purdue University. The implementation is cross-platform portable and works on Linux, Windows and Mac OS X operating systems. The MBIR software package used for ET is freely available in the form of binary executables and source codes from Dr. Bouman’s website (https://engineering.purdue.edu/~bouman/OpenMBIR/bf-em). A tutorial of MBIR can be found in the Supplementary Note.

### Test datasets

We evaluated the performance of MBIR method on both plastic embedded ET dataset and cryo-ET datasets by comparing its results with three reconstruction techniques used in the cryo-ET community, including BP and SIRT available in IMOD ^16^, and ICON ^10^. The plastic embedded ET dataset obtained from IMOD tutorial website ^16^ was originally provided for dual axes reconstruction, but we only used the first tilt series (BBa.st) in our study. Four published experimental cryo-ET datasets (EMPIAR-10037, EMPIAR-10045, EMPIAR-10064 and EMPIAR-10110) were downloaded from the public database EMPIAR ^17^. EMPIAR-10064 dataset was collected with the Volta phase plate (VPP). These tilt series were aligned based on fiducial gold markers using IMOD and then the same aligned tilt series were reconstructed by the four reconstruction techniques, respectively. The details of these datasets are summarized in Supplementary Table 2 including data type, biological sample, instrument, defocus, tilt scheme, total dose, and data collection software. In this study, the figures used for comparing the performance of different methods are contrast-normalized to avoid subjectivity of observations and to ensure the reliability of comparison.

## Acknowledgements

This work was supported in part by Showalter Faculty Scholar grant. We thank Ms. Brenda Gonzalez for her assistance in preparation of the manuscript.

## Author Contributions

R.Y. and W.J. conceived the project and designed the research, analyzed and interpreted data, drafted, revised, and completed final approval of the manuscript. S.V. and C.B. developed the algorithm and implemented the code. J.L. provided some test data. All authors reviewed the manuscript, agreed to all the contents and agreed the submission.

---

**Supplementary Fig. 1.**
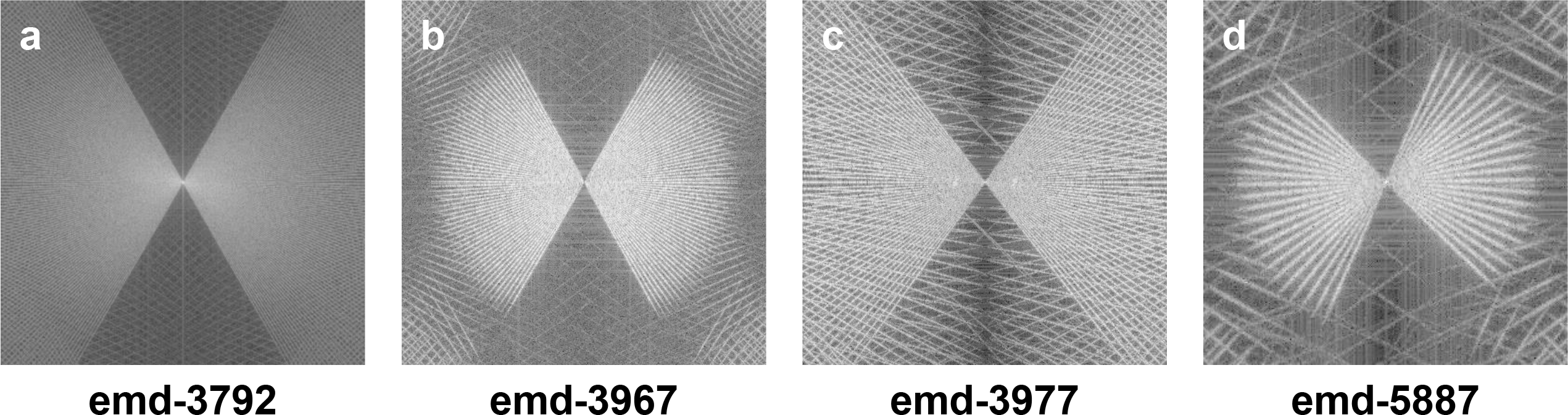
Observations of aliasing issue in the central XZ-slice power spectra of cryo-ET tomograms downloaded from EMDB. The EMDB ID is marked at the bottom of each image.

**Supplementary Fig. 2.**
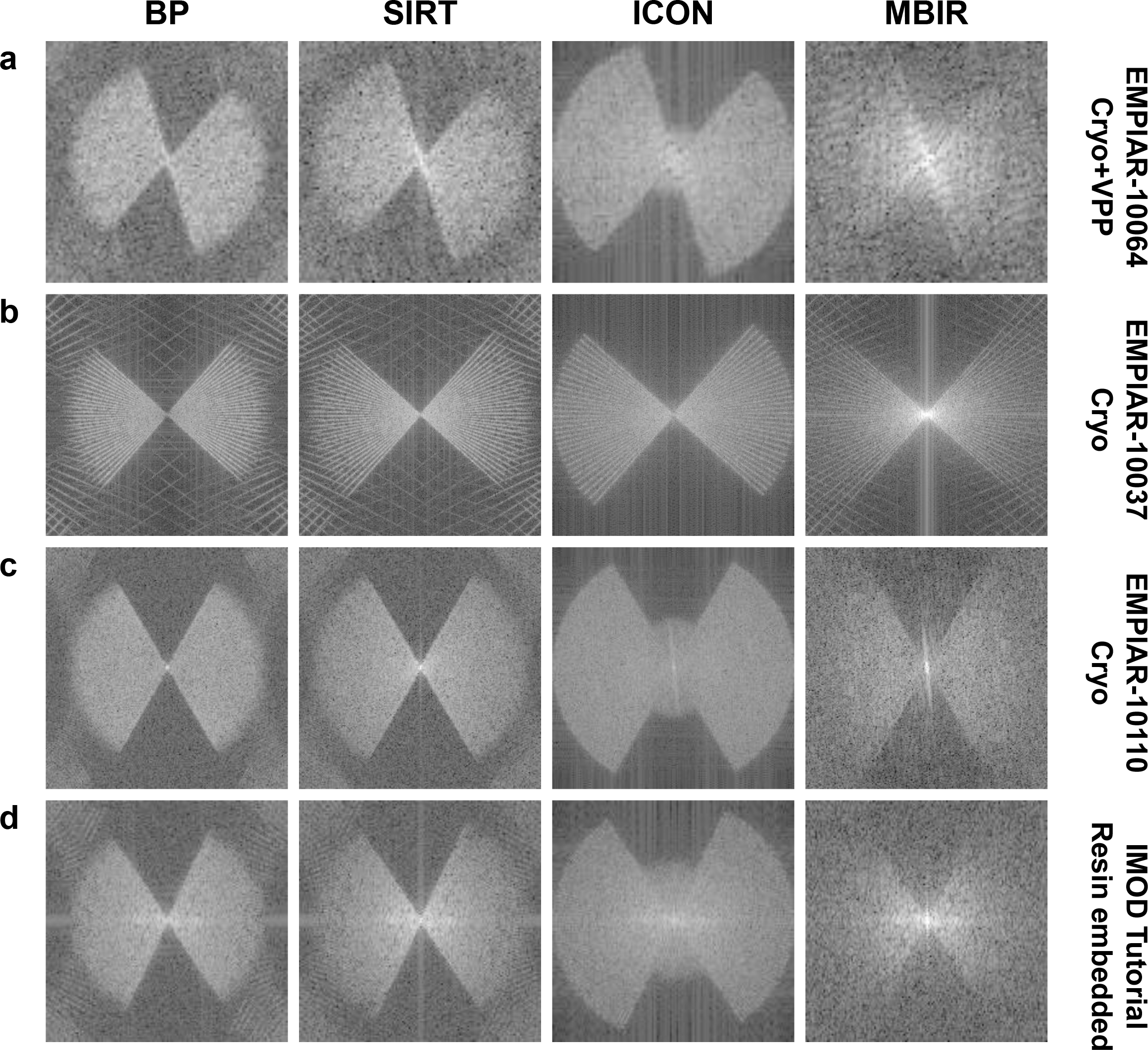
Comparison of the central XZ-slice power spectra from the tomograms reconstructed by different reconstruction techniques. Each row shows the power spectra of central XZ-slices from the same dataset using different methods. Each column shows the power spectra from the same method applied to different datasets. The data type and EMPIAR ID are denoted at the right side of each row.

**Supplementary Fig. 3.**
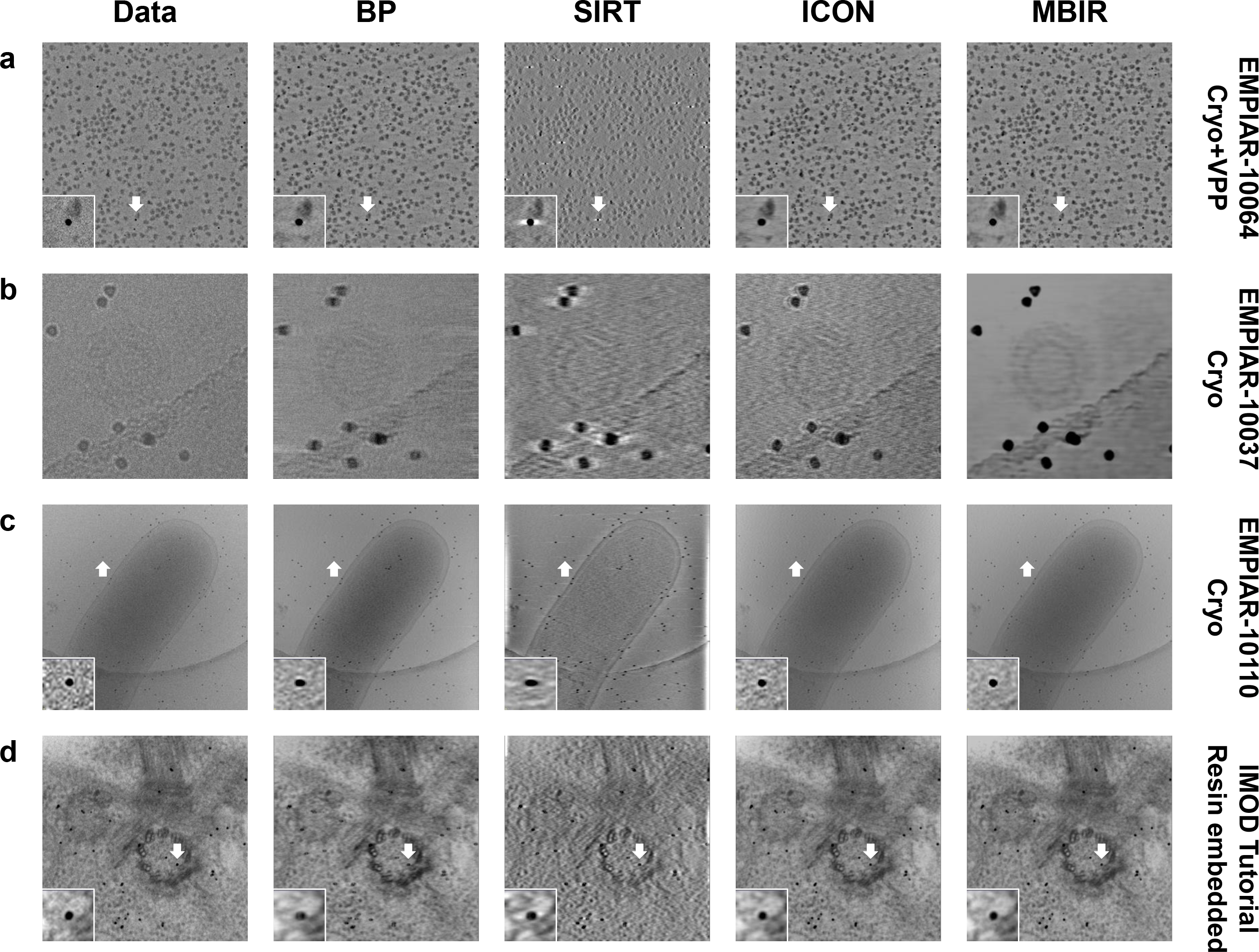
Comparison of reprojections at 0° when the tomograms are generated using different reconstruction techniques without the corresponding tilt. Each row shows the reprojections from the same dataset using different methods. Each column shows the reprojections from the same method applied to different datasets. The data type and EMPIAR ID are denoted at the right side of each row. In each image, the gold marker indicted by a white arrow is displayed with corresponding zoomed-in view.

**Supplementary Fig. 4.**
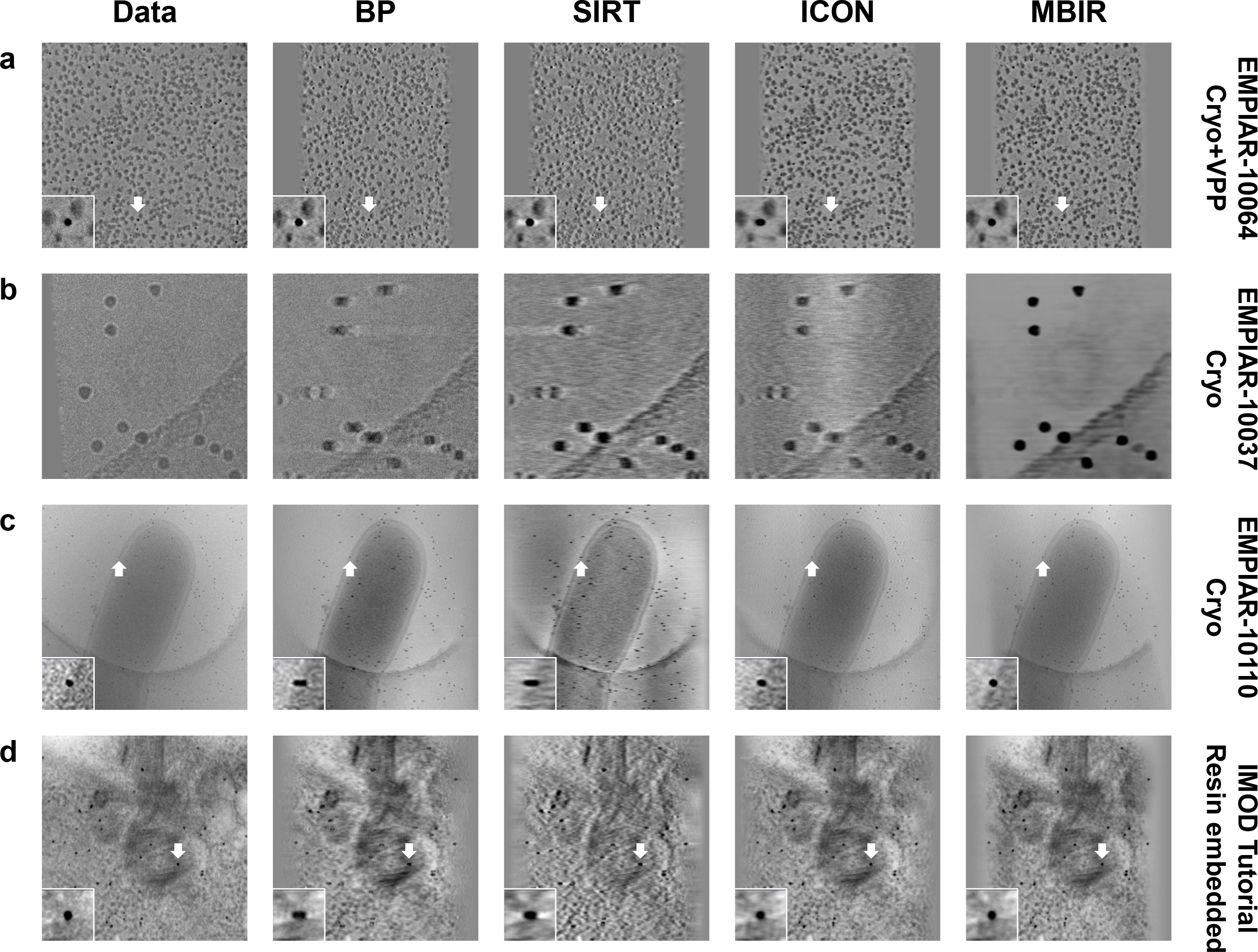
Comparison of reprojections at 45° when the tomograms are generated using different reconstruction techniques without the corresponding tilt. The details are the same as described in Supplementary Fig. 3.

**Supplementary Table 1.** Computational speeds of different reconstruction techniques. The input data size is 400 pixels × 400 pixels × 61 tilts and the output tomogram size is 400 pixels × 400 pixels × 128 pixels. The CPU model is Intel(R) Core(TM) i7-6900K @ 3.20GHz. The GPU model is NVIDIA GeForce GTX 1080.

| Method | CPU/GPU | Time (seconds) |
| --- | --- | --- |
| BP | 1 CPU core | 17 s |
| SIRT | 1 CPU core | 186 s |
| ICON | 1 GPU | 246 s |
| MBIR | 8 CPU cores | 1325 s |

**Supplementary Table 2.** A summary of datasets used in this study.

|  |  |  |  |  |  |
| --- | --- | --- | --- | --- | --- |
|  | EMPIAR-10045 | EMPIAR-10064 | EMPIAR-10037 | EMPIAR-10110 | Imod Tutorial Dataset |
| Data Type | Cryo | Cryo+VPP | Cryo | Cryo | Plastic embedded |
| Biological Sample | *S. cerevisiae* 80 S ribosome | 80S ribosome | Immature-like Rous-Sarcoma Virus Gag particles | Toxin-coregulated pilus machine | Centriole |
| Instrument | FEI Titan Krios 300KV;<br>Gatan Quantum Energy Filter;<br>Gatan K2 Summit DDD | FEI Titan Krios 300KV;<br>Gatan K2 Summit DDD | FEI Titan Krios 200KV;<br>GIF2002 Energy Filter;<br>Gatan MultiScan 795 CCD | FEI Polar 300KV;<br>Gatan Energy Filter;<br>Gatan K2 Summit DDD | Gatan Camera on TF30 |
| Defocus | 3~5 $\mu\text{m}$ | 0~200 nm | 1.5~5 $\mu\text{m}$ | ~6 $\mu\text{m}$ | Unknown |
| Tilt Scheme | -60°~+60°<br>Stepsize 3° | -60°~+60°<br>Stepsize 2°<br>First +20°~-60°,<br>then +22°~+60° | -45°~+45°<br>Stepsize 3°<br>First 0°~-45°;<br>then +3°~+45° | -60°~+60°<br>Stepsize 1° | -60°~+60°<br>Stepsize 2° |
| Total Dose ($\text{e}/\text{\AA}^2$) | 60 | 90~100 | 24~34 | 160 | Unknown |
| Data Collection Software | Serial EM | Serial EM | FEI Tomography Software | UCSF Tomography Software | Serial EM |

## Supplementary Note

### MBIR tutorial

Step 1: Download MBIR. https://engineering.purdue.edu/~bouman/OpenMBIR/bf-em/index.html

There are two Youtube application tutorials (Basic and Advanced) on MBIR website using one simulated dataset. It is recommended to watch these tutorials first. This protocol is similar to the tutorials and used to show the application of MBIR on cryo-ET dataset based on Linux system.

Step 2: Decompress the downloaded package and activate the GUI../OpenMBIR-v2.35-RHEL6-x86_64/bin/BrightFieldGui.sh

Step 3: Load your aligned tilt series as input. This software supports aligned input files stored in FEI MRC format.

Click **Select** to load your aligned tilt series. If the file suffix is not default, change **Files of type** to **All files** to access your file.

Click **Open** to load your aligned tilt series.

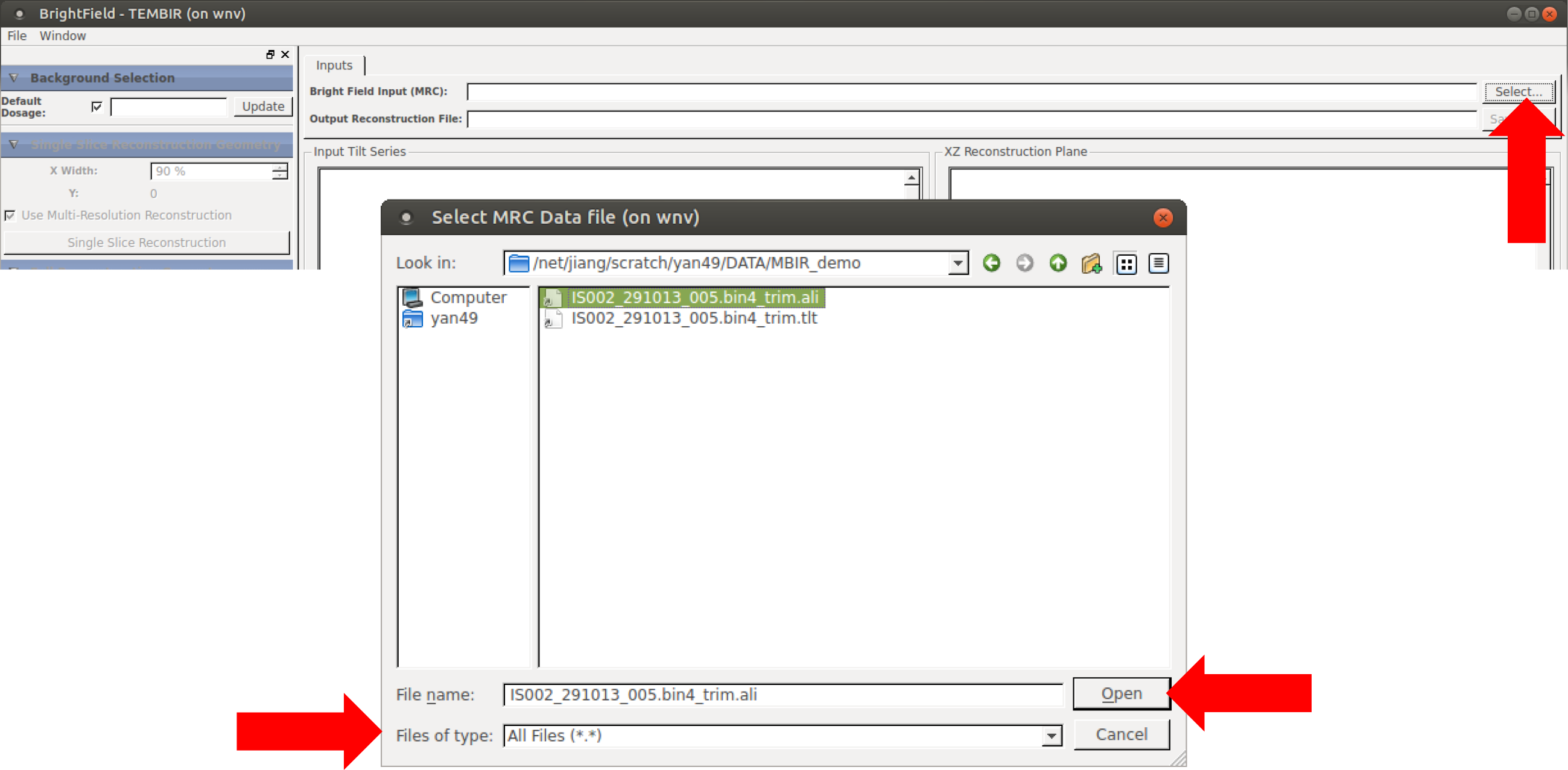

Step 4: Calculate the default dosage.

Move the red box to a background area without sample, adjust its size, then click **Update** in **Background Selection**.

Step 5: Input **Sample Thickness (nm)** in **Full Reconstruction Geometry**.

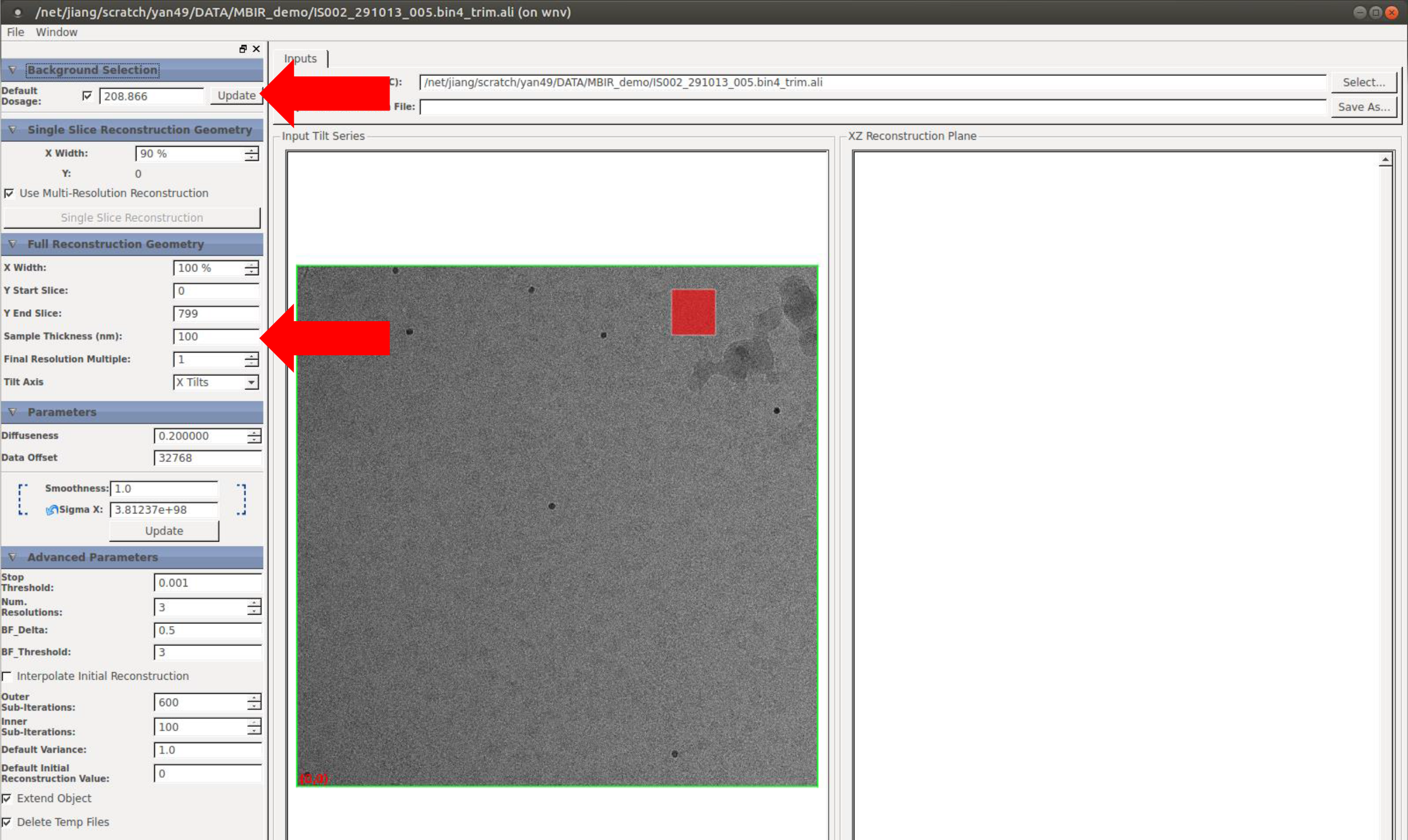

Step 6: Load your tilt angles file and input apix.

Go to upper left corner, select File **→** Load Tilt Values

In the dialog, click **Select** to load your tilt angle file. If the file suffix is not default, change **Files of type** to **All files** to access your file.

Input **Pixel Size (nm)**.

Click **OK** to complete this step.

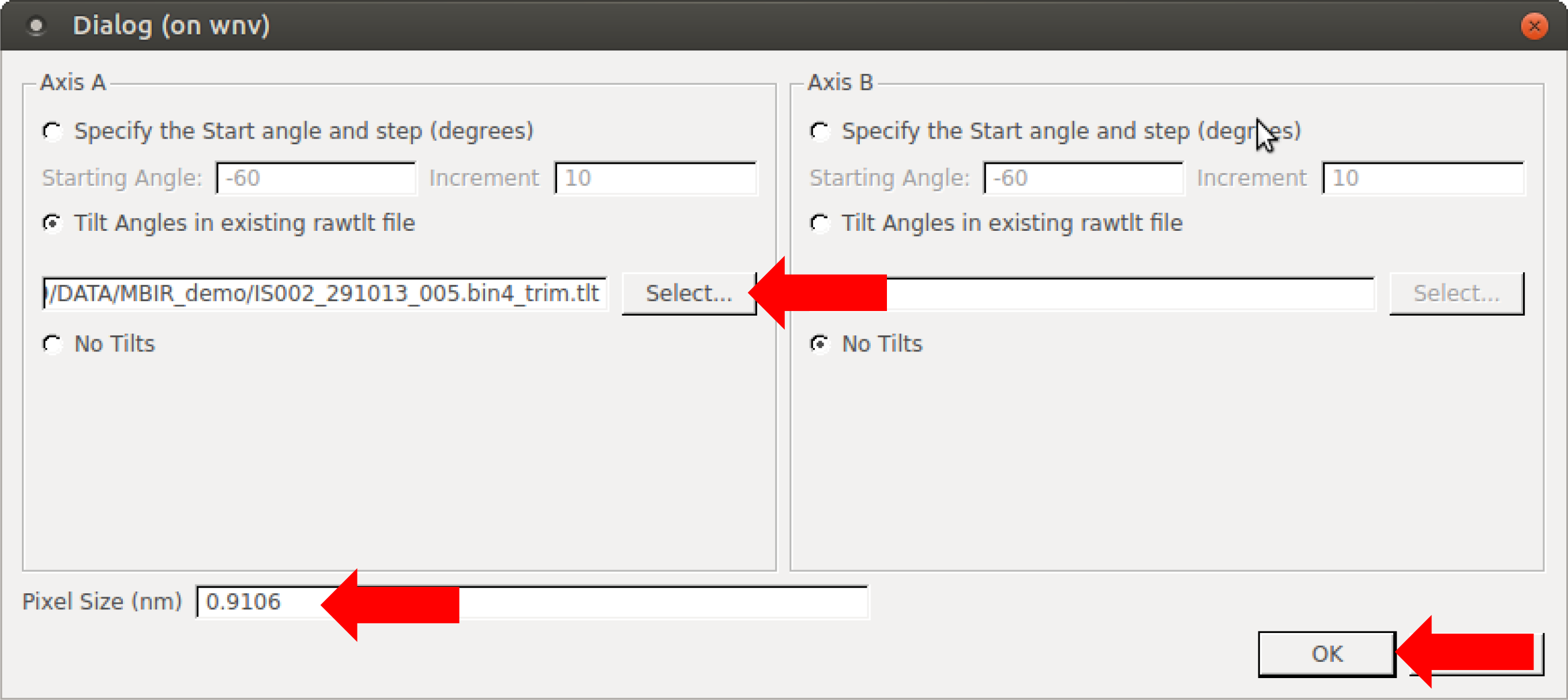

Step 7: Smoothness and diffuseness.

After loading tilt values in step 6, the software will automatically calculate **Sigma X** in **Parameters**. Wait a few seconds until **Estimating Target Gain and Sigma X Complete**
appears at the lower left corner of the GUI window. This is equivalent to click **Update** in **Parameters**. If you change the area of interest (e.g. X Width, Y start slice and Y end slice, Sample thickness), you may need to click **Update** in **Parameters** to recalculate the value of **Sigma X**.

In **Parameters**, you will see a **Sigma X** value corresponding to **smoothness** 1. Our empirical value of **smoothness** is 0.15~0.35 for cryo dataset. Here we use 0.15 for this dataset. And the default value of **diffuseness** 0.2 is good for cryo dataset.

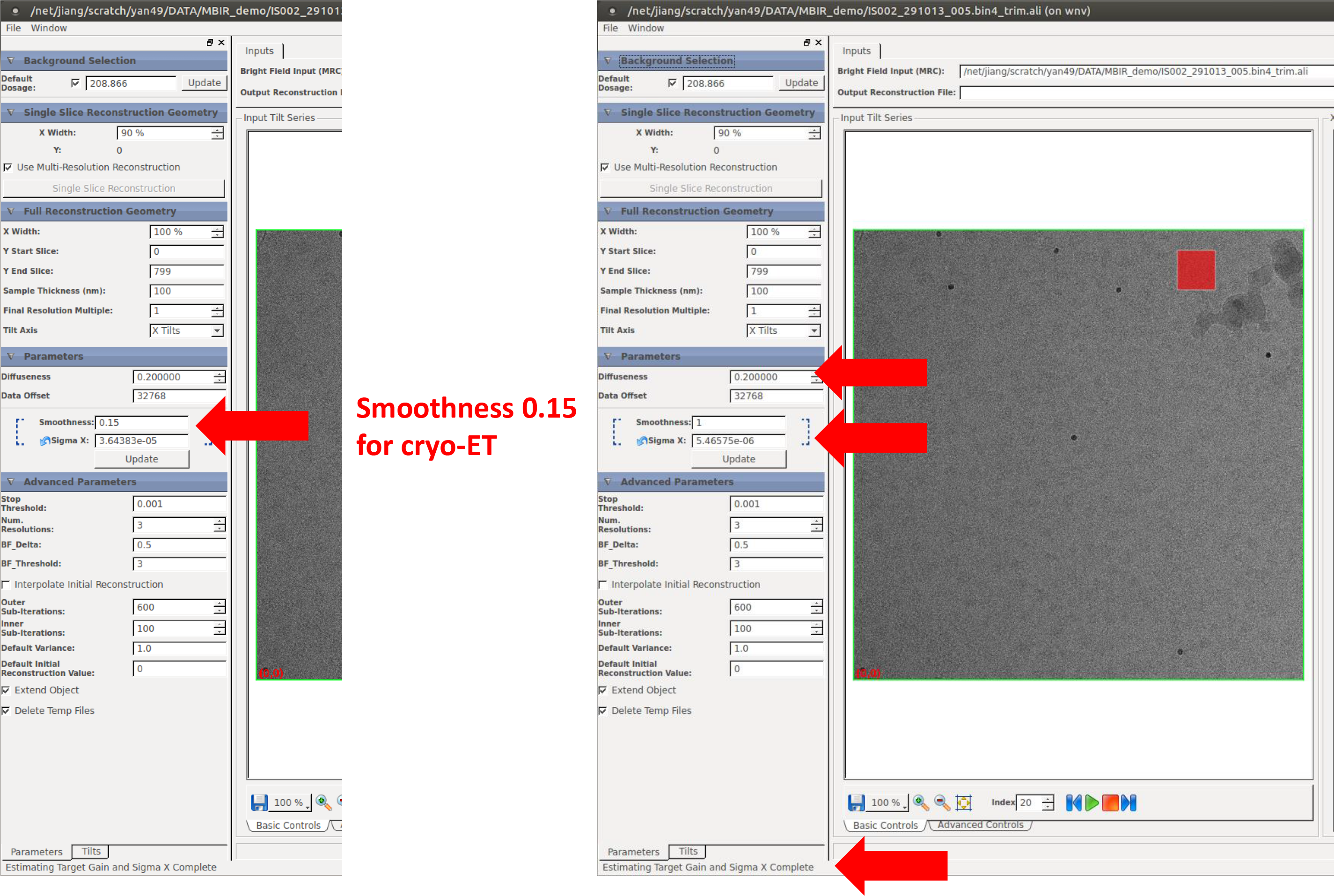

Step 8: Reconstruct your tomogram.

Click **Save As** to save your output tomogram.

Click **Reconstruct** at the lower right corner to start the reconstruction.

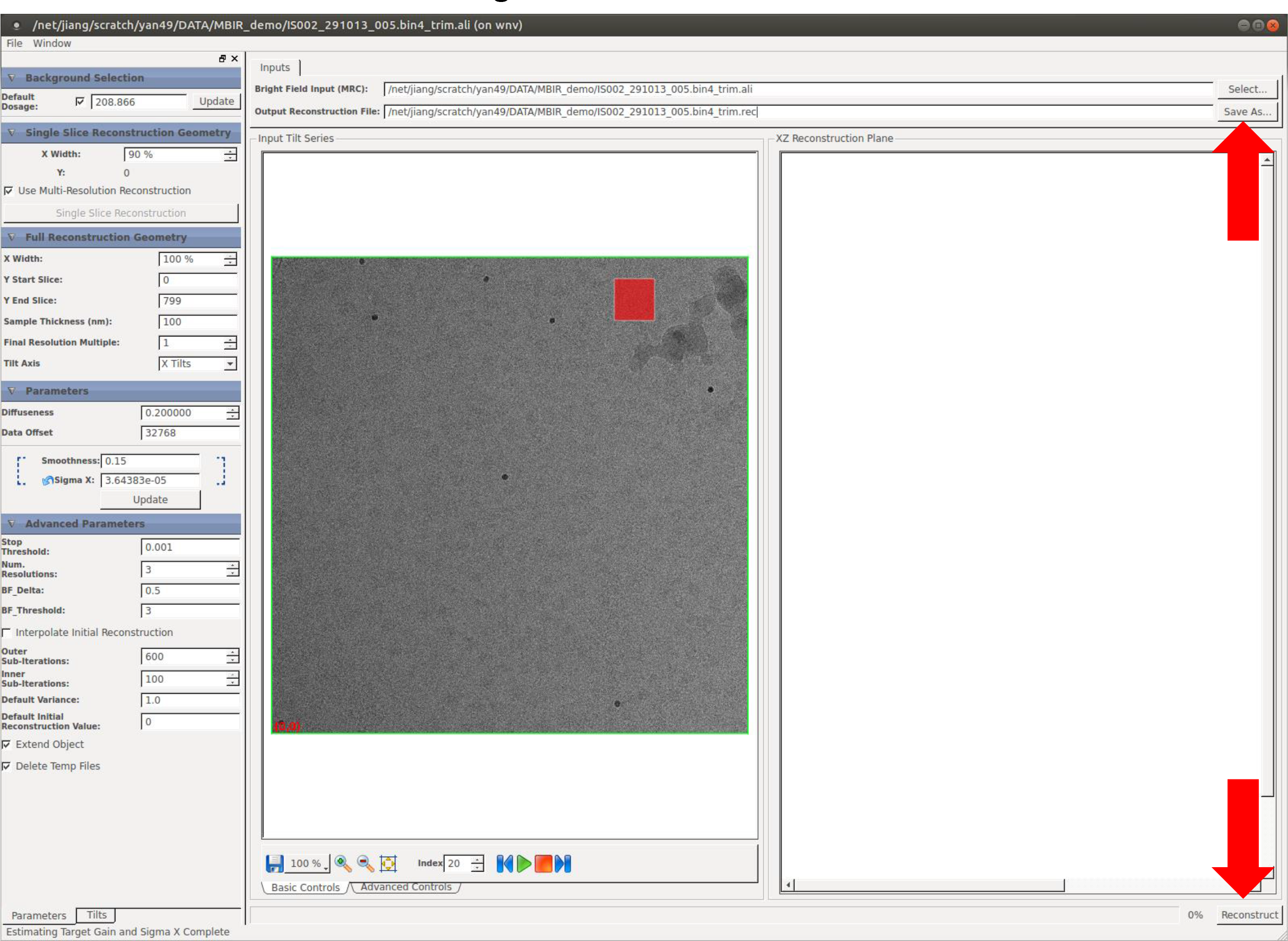

Tips:

If you do not know which smoothness is optimal for your dataset, you can either use **Single Slice Reconstruction Geometry** or reconstruct a subarea of your tilt series.

If you use **Single Slice Reconstruction Geometry**, hold SHIFT key and click somewhere on your input file, you will see a yellow line, indicating where you want to make a single XZ slice reconstruction.

Click **Single Slice Reconstruction** in **Single Slice Reconstruction Geometry** and wait several seconds. You will see a single XZ slice reconstruction shown in the right window. At this time, you can only view this single XZ slice here but not save it.

You can adjust the **smoothness** value to reconstruct different single slices and determine the optimal **smoothness** for your dataset.

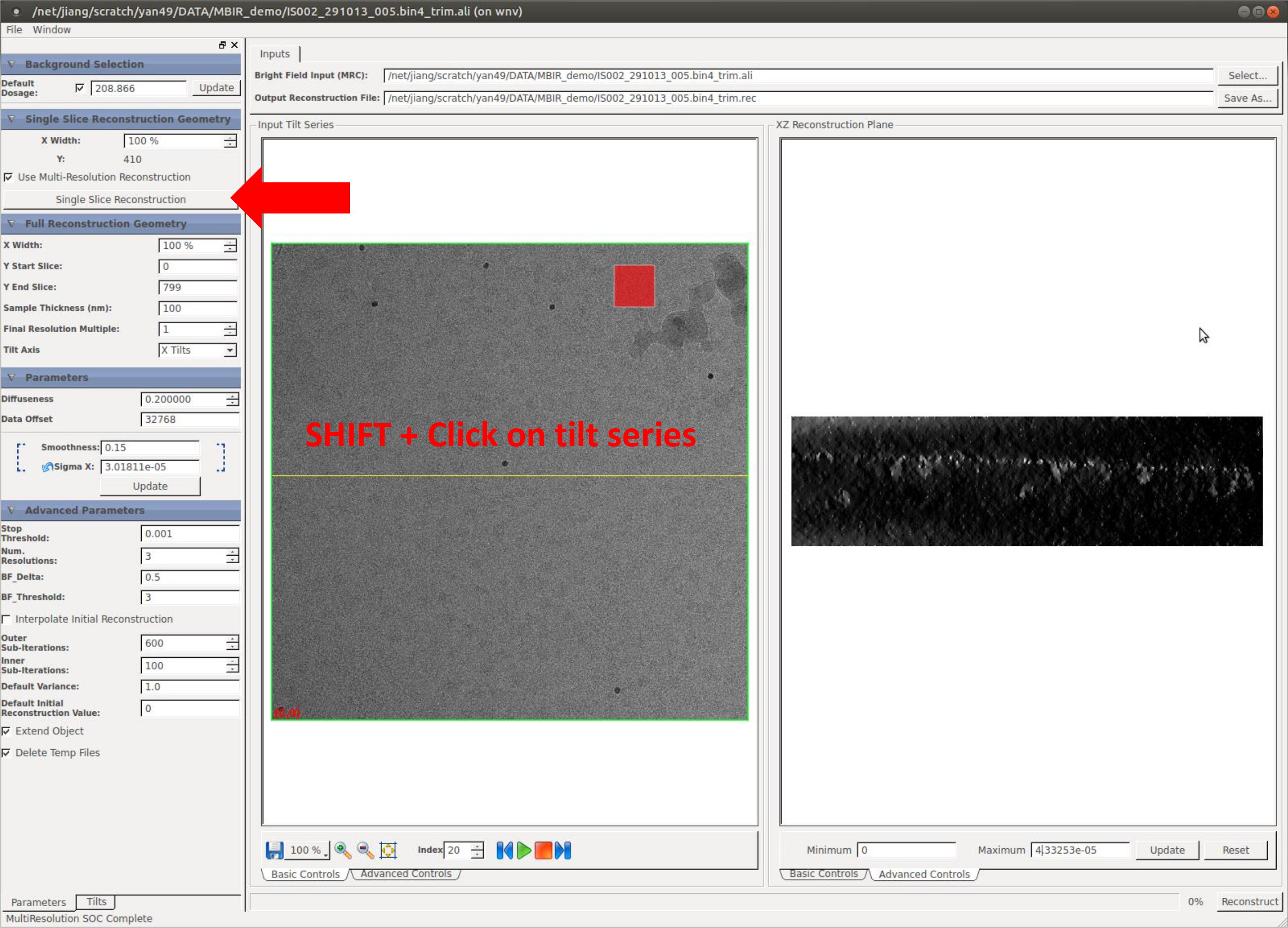

It is also recommended to reconstruct a small area to determine the optimal **smoothness**. You can adjust the top and bottom edges of the outer green box to find the **Y start slice** and **Y end slice** of your target area. When you adjust the top and bottom edges of the green box, you will see the corresponding change of the numbers in **Y start slice** and **Y end slice** in **Full Reconstruction Geometry**. However, the range of X coordinate can only be adjusted using **X Width** in **Full Reconstruction Geometry** in order to guarantee the selected area is symmetric to the tilt axis.

When you change the area of interest (e.g. X Width, Y start slice and Y end slice, Sample thickness), you may need to click **Update** in **Parameters** to recalculate the value of **Sigma X**

Click **Save As** to save your trial tomogram.

Click **Reconstruct** at the lower right corner to start the reconstruction.

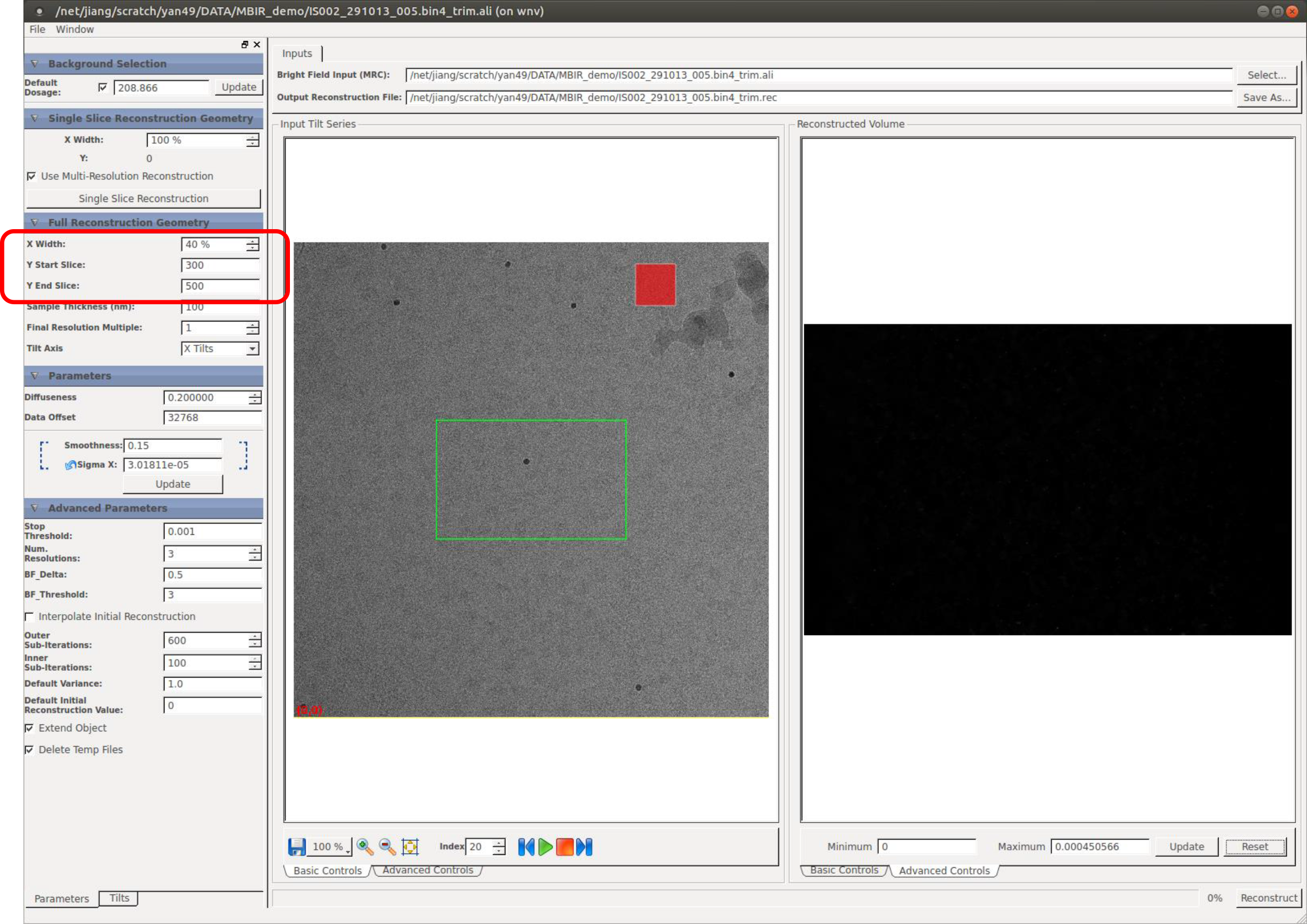

Here your trial tomogram is saved to the provided directory. You can use IMOD or other visualization software to open it and examine the quality of reconstruction.

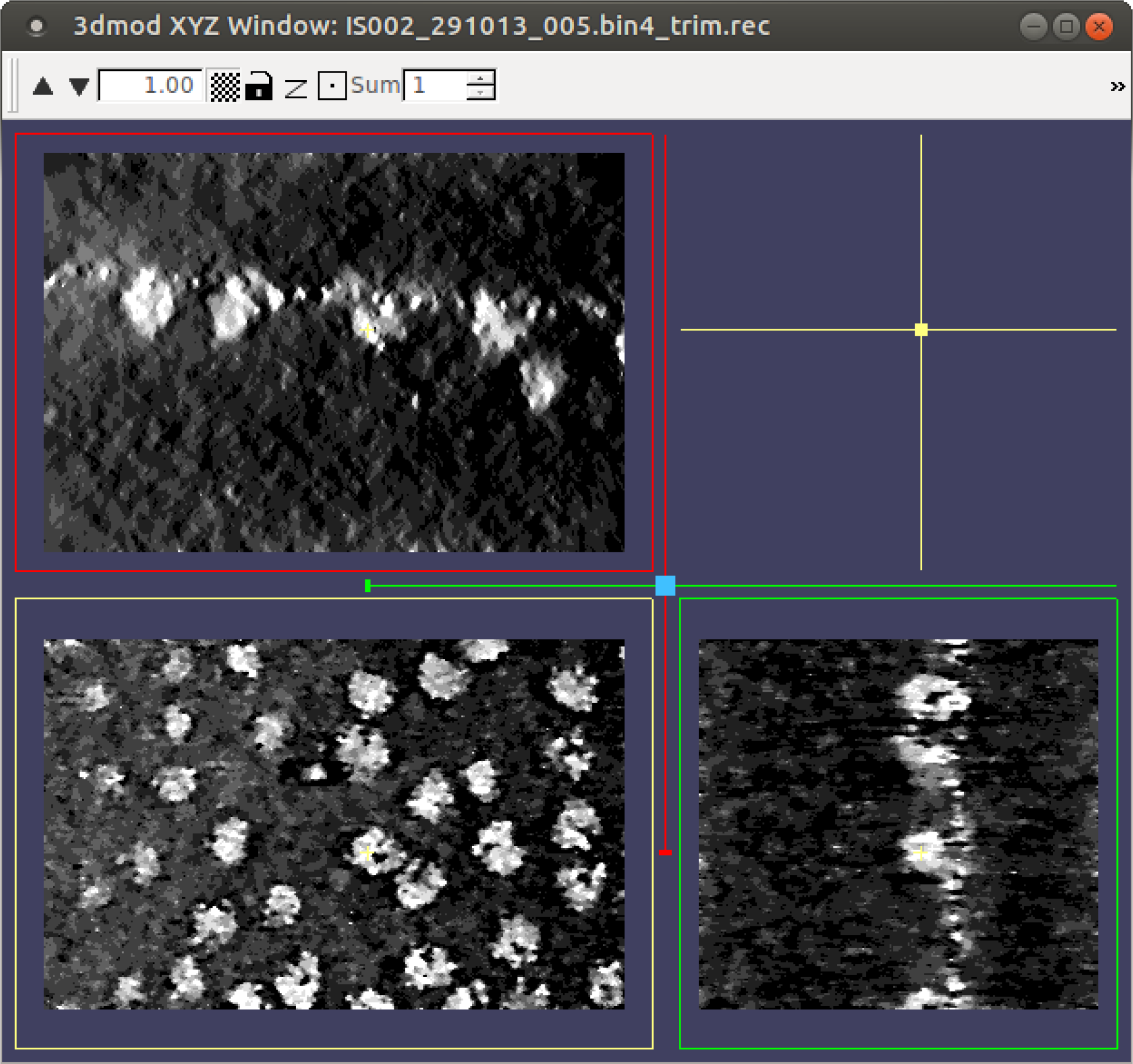

